# SCREP: Towards Single-Cell Drug Response Prediction by Pharmacogenomic Embedding Enhanced Meta-Pretraining and Few-Shot Transfer Learning

**DOI:** 10.1101/2024.04.25.591050

**Authors:** Shuang Ge, Shuqing Sun, Yiming Ren, Huan Xu, Qiang Cheng, Zhixiang Ren

## Abstract

**Objective:** Single-cell pharmacogenomic data is crucial for identifying biomarkers and understanding resistance mechanisms. However, the limited availability of those data poses significant challenges for efficient pre-training and thus hinders the generalization capability of the model. This paper aims to develop an efficient predictive model for single-cell drug response inference by leveraging knowledge from a bulk dataset.

**Methods:** To translate knowledge from bulk cell lines to single-cell analysis, this paper proposes a meta-pretraining and few-shot transfer learning framework based on pharmacogenomic embeddings. To enhance feature representation and alignment, genomic information is processed by a position-based feature extraction network to extract contextual features. Simultaneously, a graph-aware Transformer is developed to capture interatomic relations for drug information representation. Underlying this framework, key drug action pathways can be identified at the cellular level through the proposed gene gradient attribution algorithm.

**Results:** This model integrates drug response data from 223 drugs across 14 tissues for meta pre-training, followed by transfer and testing on seven single-cell datasets. In comparison to other models, the proposed framework achieves an average accuracy increase of 4.58% for pre-trained drugs and demonstrates a 20% improvement in generalization performance for previously unseen drugs. Case studies further illustrate its capability to differentiate between resistance genes.

**Conclusion:** The proposed framework exhibits enhanced generalization capabilities in predicting cellular responses across multiple drugs, including unseen drugs. This model serves as an effective tool for investigating drug action pathways and elucidating resistance mechanisms from single-cell insights, with significant implications for guiding clinical medication practices.

## 1 Introduction

From a clinical perspective, accurately evaluating drug response in cancer patients is vital for effective treatment. Studies have shown that alterations in genomic profiles significantly influence the success of cancer therapies[1, 2] and can be regarded as biomarkers to identify populations sensitive to treatment. The heterogeneity of genomic profiles poses substantial challenges in accurately predicting responses to personalized anti-cancer therapies[3]. In this context, single-cell RNA sequencing (scRNA-seq) analysis has emerged as a powerful tool, providing novel insights into the cellular composition of tumors[4] and enabling the identification of distinct cellular subpopulations that contribute to resistance[5, 6]. The analysis of scRNA-seq holds great promise for enhancing drug response prediction, tailoring therapies to individual patients, and facilitating personalized oncology[7].

However, studies on drug responses at the single-cell level (sc-level) currently encounter several challenges. These difficulties primarily stem from the limited pharmacogenomic data associated with scRNA-seq datasets and the inherent complexity of data analysis. However, numerous studies have successfully used drug response profiling in bulk cell lines. These cell lines have been developed to represent a diverse array of cancer types originating from primary tumors, effectively preserving tissue characteristics and the immune microenvironment [3]. Numerous studies have demonstrated that pharmacogenomic profiling of bulk cell lines has become a platform for biomarker discovery and for predicting in vitro drug responses to various targeted therapies [8, 9]. In this context, the potential of pre-training on bulk cell lines and transferring bulk knowledge to single-cell level analyses emerges as a promising strategy to address these challenges. A few recent studies, such as scDEAL[10], and SCAD[11] attempt to leverage publicly available drug-cell line sensitivity profiles to predict drug response at the cellular level. However, prior studies have predominantly used transcriptomic information to pre-train drug-specific models, which lack pharmacogenomic insights for other drugs and thus hinder model generalization.

Leveraging both pharmaceutical and genomic information during pre-training can provide a wealth of domain knowledge, thereby enhancing the generalization ability for predicting responses across different drug-cell line. Prior work has suggested that structurally similar molecules exhibit analogous properties, especially when structural similarities indicate shared interactions with biological targets[12]. By integrating molecular information during the training phase, the model can effectively learn the interactions between drugs and genomes. It is not restricted to a specific drug. This methodology not only improves the accuracy of predictions but also expands their applicability across various therapeutic contexts. This, in turn, facilitates drug repurposing [13] and supports the identification of novel therapeutic targets [14].

We innovatively combined task-specific multimodal representations with meta pretraining to develop a transfer learning framework called SCREP, aimed at leveraging extensive bulk knowledge to enhance modeling of single-cell drug response prediction. To comprehensively extract the knowledge embedded in the compounds, we employ a graph-aware Transformer network to model the interrelations among atoms. Subsequently, we employ a position-based neural network to process genomic information organized in a specific sequence, thereby facilitating the extraction of rich contextual features. Experimental results demonstrate that our model generalizes effectively to previously unseen drugs, while case studies further highlight its interpretability and practical applicability. In summary, our primary contributions are as follows:

- This paper presents a novel meta pre-training and adaptive transfer framework that integrates representations of molecular to predict the response of multiple drugs at cellular level.
- This paper proposes a few-shot adaptive distribution alignment strategy for cross-domain migration, facilitating task-specific few-shot fine-tuning while minimizing sample size requirements.
- A graph-aware Transformer network is designed to capture complex atomic interactions within drug molecules, and a convolutional neural network is developed to extract location-based genomic features.
- This paper evaluates the generalization capabilities of the proposed model on over 100 drugs unseen during the pre-training phase, achieving an average classification F1 score of 87.06%.

The rest of this paper is organized as follows. In Section 2, we briefly reviewed the existing work in sc-level drug response prediction that translate from bulk-level, mainly introduce methods about deep transfer learning. In Section 3, we describe the details of the data, model, strategy, implementations and evaluations. In Section 4, we evaluate the effectiveness of transferring pharmacogenomic knowledge from bulk RNA sequencing (bulk RNA-seq) to scRNA-seq. We perform an ablation study to understand the impacts of knowledge transfer from bulk RNA-seq to scRNA-seq. Additionally, we explore the model’s performance on unseen drugs and conduct a case study to explore the practical applications of the model. In Section 5, we further discuss our results and discovery, and summarize in Section 6.

### Problem

The limited availability of single-cell drug response data poses significant challenges for conducting large-scale pre-training, thereby impacting the model’s generalization capabilities.

### What is Already Known

Although numerous prior studies have utilized bulk data from specific drugs for pre-training and subsequent application to single-cell contexts, a significant limitation of these approaches lies in their scalability to the other drugs and generalizability to previously unseen drugs.

### What this Paper Adds

This study aims to develop a general framework for meta-pretraining and few-shot transfer learning using drugs as meta-pretraining tasks. It seeks to fully leverage prior knowledge from a wide range of drug compounds and heterogeneous bulk genomic data to enhance generalization in predicting drug responses at the single-cell level through novel feature representations and modeling methods.

## 2 Related work

### 2.1 Sc-level drug response prediction

Numerous studies have employed high-throughput drug screening on cancer cell lines to determine their responsiveness to various drugs. However, most of these studies confined to the cell-line level and often overlooked the impact of cell subpopulation heterogeneity on drug sensitivity. The emergence of sc-level drug response prediction, which aims to assess drug response at the cellular level, presents an opportunity to explore tumor heterogeneity from new insights. While there have been efforts to leverage deep learning networks for sc-level drug response prediction by translating knowledge from drug-cell line interactions, the optimization of these model structures tends to be drug-specific[15], e.g. scDEALL[10] and SCAD[11]. These methods require substantial amounts of target data and extensive parameter tuning for each drug to reach an optimal state suitable for transfer learning. This requirement may limit their applicability across a wide range of drugs or facilitate high-throughput drug screening[15].

### 2.2 Deep transfer learning method

Unlike traditional machine learning methods, deep transfer learning can often handle situations where the distributions of source and target data differ significantly. Techniques such as adversarial discriminative domain adaptation (ADDA)[16] and domain-adaptive neural network (DaNN)[17] have been applied to predict sc-level drug responses under bulk-level pre-training. However, these methods face inherent challenges that make it difficult to optimize networks and apply to previously unobserved drugs. On the other hand, these networks often require a large number of samples to align feature distributions, which may not always be feasible. In contrast, few-shot learning strategies are designed to learn efficiently from a minimal number of examples. By learning to recognize patterns from only a few instances, the model can understand the heterogeneity of the data, potentially enhancing their performance on unseen data or tasks, making them more versatile and efficient in data-scarce scenarios.

## 3 Materials and methods

### 3.1 Data collection and processing

#### 3.1.1 High-throughput drug responses datasets

The bulk pharmacogenomic datasets are collected from GDSC1000 project[3], which systematically tests cell line responses by high-throughput drug screening and provide half-maximal inhibitory concentration (IC50) values as metrics for drug sensitivity. Moleculars selected for screening are mainly anti-cancer therapeutics, covering both targeted agents and cytotoxic chemotherapies. Following the approaches of previous studies such as SCAD[11], GraphDRP[18] and GraTransDRP[19], we focused on 223 drugs applied to 1018 cell lines, with drug response values in terms of IC50 normalized within a range of (0,1). In addition, the GDSC bulk RNA-seq data was downloaded from https://www.cancerrxgene.org/downloads/bulk download. To assess the robustness of our methodology across various high-throughput drug response datasets, we additionally gathered pharmacogenomic data from the Cancer Cell Line Encyclopedia (CCLE) to serve as a pre-trained dataset. We gathered genomic profiles from the CCLE dataset [3] and combined them with drug response data from the PRISM Repurposing dataset. The PRISM dataset includes 1,448 compounds tested across 499 cell lines [26]. This dataset is publicly accessible through the Broad DepMap Portal^1^.

#### 3.1.2 scRNA-seq datasets

For post-treatment data, we collected dataset 1-4 (the first five datasets in the Table 1) from National Center for Biotechnology Information’s Gene Expression Omnibus (GEO)^2^. Dataset 1 and 2[20] characterizes resistance and metastasis during tumor evolution in oral squamous cell carcinoma (OSCC), including heterogeneous cell subpopulations, posing a challenge for computational tools in predicting Cisplatins sensitivity. Dataset 3[21] uses single cell sequencing data from non-small cell lung cancer to explore the mechanisms leading to tyrosine kinase inhibitor (TKIs) resistance, in particular the characteristics of resistant cell populations and their differences from drug-sensitive cells. Dataset 4[22] uses acute myeloid leukemia (AML) cells for resistance screening and identified the key role of specific regulators in epigenetic resistance through CRISPR-Cas9 screening experiments. For prior-treatment data, we obtained the pan-cancer dataset 5 and 6 (the sixth dataset in Table 1) from Kinker et al.[23] available from the Broad Institute’s single cell portal (SCP542). In their study, Kinker et al. demonstrated that EpiSen-high and EpiSen-low cells, derived from the same cell lines, exhibit differential responses to a variety of drugs[23]. Building on this foundation[11], we adopted a threshold using the 10th percentile EpiSen score to define the sensitivity of each cell to all drugs. Dataset 7[24] features pre-labeled populations of sensitive and resistant cells to Afatinib, identified through lineage tracing. A detailed description of each dataset is provided in Table 1. The scRNA-seq data and metadata files were all collected from previous studies. The above datasets are labeled data and experimentally validated, which are suitable for the validation of computational methods. These datasets specifically focus on the study of tumor heterogeneity and resistance mechanisms, which present significant challenges for computational approaches. Moreover, the effectiveness of computational methods can be used as an auxiliary tool in experimental science, thereby facilitating scientific discovery. Following the preprocessing strategy used in scBERT[25], we utilized genes from the PanglaoDB dataset as a reference to unify the dimensions across both bulk RNA-seq and all scRNA-seq datasets. The PanglaoDB dataset[26], which includes 16,906 genetic symbols, was downloaded from the PanglaoDB website^3^. To ensure consistency in gene nomenclature, the test dataset of SC transcriptomic were pre-processed by revising gene symbols according to the NCBI Gene database^4^. This step is essential to ensure that the gene names in the dataset correspond to the symbols in PanglaoDBDB, allowing us to obtain gene expression profiles that are dimensionally consistent.

**Table 1.** Summary of the seven scRNA-seq datasets. All datasets are public avaliable from National Center for Biotechnology Information’s Gene Expression Omnibus (GEO)

|  | GEO access | Drug | Cell line | Cancer type | No. Res | No. Sens | N.A. | Author |
| --- | --- | --- | --- | --- | --- | --- | --- | --- |
| 1 | GSE117872_HN120 | Cisplatin | OSCC | Oral squamous cell carcinomas | 172 | 346 | / | Sharma, et al[20] |
| 2 | GSE117872_HN137 | Cisplatin | OSCC | Oral squamous cell carcinomas | 150 | 388 | / | Sharma, et al[20] |
| 3 | GSE149383 | Erlotinib | PC9 | Lung cancer | 617 | 849 | / | Aissa, et al.[21] |
| 4 | GSE110894 | I-BET-762 | leukaemic cells | Acute myeloid leukemia | 670 | 719 | / | Bell, et al[22] |
| 5 | GSE157220 | Gefitinib, Vorinostat, AR-42 | SCC47 | Head and Neck Cancer | 162 | 162 | 472 | Kinker et.al[23] |
| 6 | GSE157220 | NVP-TAE684, Afatinib, Sorafenib | JHU006 | Head and Neck Cancer | 81 | 81 | 259 | Kinker et.al[23] |
| 7 | GSE228154 | Afatinib | MDA-MB-468 cells | metastatic breast cancer | 665 | 846 | / | J. M. McFarland[24] |

We performed quality control and preprocessing of the scRNA-seq data using the Python package SCANPY[27]. This involved filtering out cells that exhibited fewer than 200 genes, and genes that appeared in fewer than three cells. Then, cells with more than 10% expression of mitochondrial genes, which could suggest compromised cell integrity, were excluded. The count matrix was then normalized using the shifted logarithm method[28] to stabilize the variance across cells. Following normalization, expression values were scaled using the MinMaxScaler function from the preprocessing module in the sklearn package, which is suitable for preparing data for deep learning networks. Different from the bulk dataset, all sc datasets were labeled as sensitivity or resistance.

#### 3.1.3 Unseen drug datasets

The unseen drug datasets, collected from GSE157220, include 108 drugs that are not present in the GDSC dataset. To maintain consistency in data processing and analysis, we applied the same preprocessing pipeline as used for dataset 5 and 6.

#### 3.1.4 Drug molecular representations

For chemical information about drugs, the notation of each molecular were downloaded from PubChem[29] in SMILES[30] format, based on the drug names and PubChem IDs provided by the GDSC. These chemical structure information was preprocessed using RDKit[31], an open-source cheminformatics toolkit, to construct molecular graphs where atoms serve as nodes and chemical bonds as edges. Atom features were constructed using five attributes: atomic symbol, atom degree (calculated from the number of bound neighbors including hydrogen), total number of hydrogens, implicit valence of the atom, and aromaticity. These features were encoded into an one-hot multidimensional feature vector following the translation rule for each SMILES string. This approach facilitates a detailed characterization of the molecular structure, which is crucial for the analysis of drug properties and interactions.

### 3.2 SCREP framework

#### 3.2.1 Overview

In the pre-training phase, we use all available bulk cell line-drug response datasets to develop a “learn how to learn” network. This network is designed to quickly adapt when presented with a few additional sc samples. The model is specifically trained to identify commonalities and distinctions between tasks, utilizing this information to facilitate new tasks effectively. We adopted a meta pretraining and transfer learning strategy for few-shot learning scenarios, and tailored our framework to better suit the requirements of sc drug response prediction. This modification ensures that the model can flexibly handle cross-domain challenges and label inconsistency issues. An overview of the SCREP framework is shown in Figure 1(a) and (b).

**Fig. 1.**
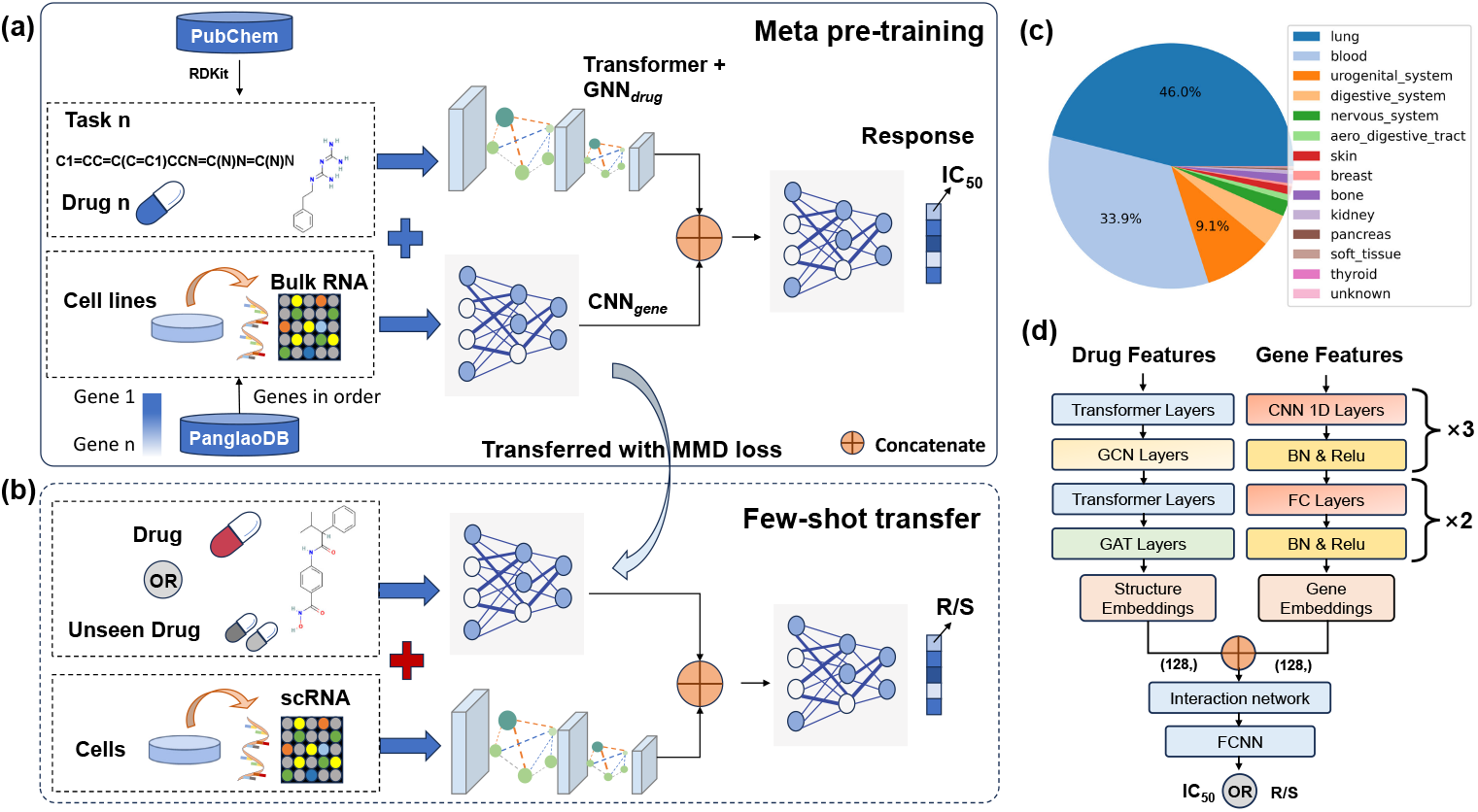
Description of the SCREP workflow. **a.** An overview of our proposed framework, including data collection and pre-processing, task-specific representation, and three modules for feature extraction and feature interaction. **b**. An overview of the few-shot transfer framework, including task-specific few-shot fine-tuning. **c**. Dataset statistic shows the distribution of bulk datasets, including 14 tissues with 58 subtissues (Figure S1). **d**. The details of each modules, including Transformer-*GNN*_*drug*_, *CNN*_*gene*_ and task-specific interaction network.

#### 3.2.2 Meta pre-training design

A meta pre-training framework is developed where each drug in the bulk dataset is treated as a distinct meta pre-training task, resulting in a total of 223 tasks corresponding to 223 drugs. For each task *T*, we further categorize the cell lines into 14 tissues as depicted in Figure 1(c), with each tissue representing a set of learning samples. During each pre-training iteration, we randomly select a dataset *S* corresponding to one drug and then sample a subset of 8 tissues *S*_*i*_ to create a smaller dataset. Our selection of eight tissues is based on previous studies, which indicates that an excessively large subset can lead to increased computational complexity, whereas a very small subset may not provide adequate information for effective generalization across various tasks. By choosing eight tissues, we acheive a balance between ensuring sufficient samples for learning within each batch and avoiding overly large sample sizes that could increase computational complexity or exceed memory constraints. This subset *S*_*i*_ is subsequently divided into two non-overlapping groups, each containing four tissues. These groups are designated as cell lines *Q* and cell lines *K*, which serve distinct roles in the training process. For the loss function, we employ the Mean-Square Error (MSE) function *L* as the pre-training loss,

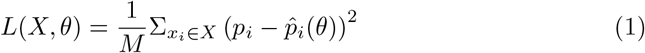

The loss function *L* is defined as a function of the sample set *X*, and the model parameters *θ*, where *p*_*i*_ represents the continuous label for sample *i* and *M* is the total number of samples in *X*.

We first perform one-step gradient descent on the subset *Q*, using it to compute a preliminary update to the parameters *θ*. Following this initial step, we then focus on optimizing the parameters further by seeking a better solution for subset *K*. This is achieved through iterative refinement, where we apply the Adam gradient descent algorithm[32] to minimize the regression loss. For each training iteration, we aim to find the optimal parameters *θ* that result in a smaller regression loss. This involves updating the parameters over 100 iterations specifically on subset *K*. The entire pre-training phase comprises 1000 training iterations, allowing us to iteratively refine the model using 1000 paired subsets *Q* and subsets *K*, respectively. For each *S*, a loss function is defined as follows:

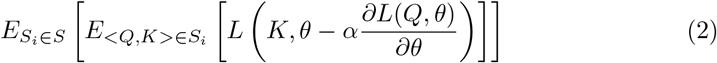

Where *L* is MSE loss as described in equation(1). For each training iteration, we seek the parameters *θ* that can achieve a smaller regression error on subset *K* after only one iteration of the gradient descent performed on subset *Q*.

#### 3.2.3 Task-specific Pharmacogenomic Embedding

##### Chemical embedding

For each drug *d*, a graph-transformer network is designed to learn the representations of molecular chemical structure (Figure 1(d)). The input to this network consists of graph features represented as 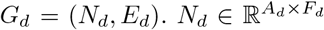 is the node matrix of the graph, where each node corresponds to an atom within the molecular structure. 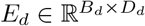 represents the adjacency matrix of edges, detailing the connections between the nodes, i.e., the chemical bonds. Where *A*_*d*_ denotes the number of atoms in the drug molecule, *B*_*d*_ represents the number of chemical bonds connecting these atoms. *F*_*d*_ corresponds to the feature embedding of the atomic properties. *D*_*d*_ denotes the bond endpoints, which are consistently established at a value of 2, given that each bond connects two atoms. The combination of a Transformer architecture with a Graph Neural Network (termed Transformer-*GNN*_*drug*_) effectively represents the interatomic relationships and contextual structural characteristics of compounds. This approach leverages the Transformer’s capability to handle long-range dependencies within sequential data, enhancing the representation of linear sequences such as SMILES strings. Simultaneously, the Graph Neural Network component can capture the intricate inter-connectivity of nodes (atoms), thus reflecting the actual spatial structures and bonding relationships in molecular.

##### Cell line gene expression embedding

The gene feature extraction network is implemented as an end-to-end convolutional neural network (CNN) without manual feature extraction. This approach allows exploiting the high-dimensional feature representations of deep neural networks, ensuring that the true genetic signature is fully preserved. At the same time, gene expression is reorganized in accordance with the gene symbol order of the PanglaoDB dataset, ensuring that the gene arrangement remains consistent across all datasets. As a result, the genomic features of aligned gene sequences contain positional relationships between contexts. Convolutional neural network (CNN) can effectively capture the semantic relationship between gene contexts due to its ability to extract local patterns through convolutional filters. Gene alignment avoids semantic mismatches between samples caused by the disruption of gene order.

#### 3.2.4 Few-shot transfer learning design

In the few-shot transfer phase, each sc dataset is treated as a new task with only a few additional samples available. Due to the distribution gap between bulk RNA-seq and scRNA-seq,the pre-trained weights is used as initialization parameters and refine all parameters during fine-tuning. For each task, *n*-shot(*n* = 1, 5, 10) with 10-query samples is preformed to update the parameters, and the remaining samples are used for testing. One iteration of gradient descent is computed on the shot set, followed by 100 iterations of updating the parameters on the query samples. This process is repeated 10 times for each *k*. Then the average and standard deviation of the prediction performance is calculated across all these repetitions (Figure 3 (c), (d)). The binary cross entropy (BCE) loss is employed with m (*m* ∈ 0 ∼ 1000) iterations, activating an early stopping mechanism if the area under the curve (AUC) of the query sets reaches 0.99. In addition, the maximum mean discrepancy (MMD)[33] loss is we incorporate to align the hidden spatial feature distribution of the genes:

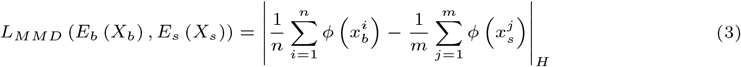

MMD is a measure used to determine the difference between the average embeddings of two distributions within a high-dimensional feature space known as the reproducing kernel hillbert space (RKHS) denoted by *ϕ*(.). Specifically, 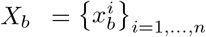 and 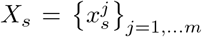 represent the embeddings of gene features from bulk cell lines *n* and single cells *m*, respectively. The intuition is that the identical distributions *E*_*s*_ and *E*_*b*_ will have similar average representations when the MMD values are close to zero. As a result, the total loss for our model is computed as the sum of the BCE loss and the MMD loss:

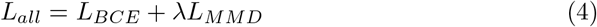

Where *λ* is introduced to balance the influence of the alignment loss. After *m* iterations of gradient descent, the few-shot parameter *θ* is adapted to the new samples:

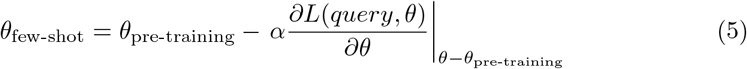

The pseudo-codes of SCREP are illustrated in Algorithm 1.

#### 3.2.5 Model training protocols

Our deep learning model constructs an end-to-end pharmacogenomic interaction model. By extracting long-range sequence features of the transcriptome and graph features of the chemical information, the model can automatically extract interaction features to distinguish pre-existing resistant cell lines under specific drugs. As depicted in Figure 1, in the meta pre-training phase, SCREP requires three steps to obtain the prediction results. First, drug features are extracted by converting the drug SMILES into a molecular graph, which is then fed into the Transformer-*GNN*_*drug*_ module. At the same time, the expression profiles of genes are processed through the *CNN*_*gene*_ module to extract features from the gene expression. Subsequently, the outputs from both the drug and gene modules, standardized to 128 dimensions, and concatenated into a nonlinear feature convergence network designed to predict the mutual relationships, ultimately estimating the IC50 value. Within the interaction model, the hidden layers are configured with 1024 and 128 neurons, respectively. The *CNN*_*gene*_ module includes three convolutional kernels of size 8, followed by two fully connected layers that map to a 128-dimensional hidden space. The Transformer-*GNN*_*drug*_ module integrates the capabilities of graph convolutional networks (GCN)[34] and graph attention networks (GAT)[35], each following a transformer layer that serves as an invariant dimension feature extractor. During the few-shot transfer phase, the model retains the same architecture as the pre-trained model but outputs predictions using binary drug sensitivity or resistance labels. Therefore, the pre-trained weights is loaded up to the last fully-connected layer for task-specific fine-tuning.

##### Algorithm 1

The pre-training and fine-tunning of our proposed framework.

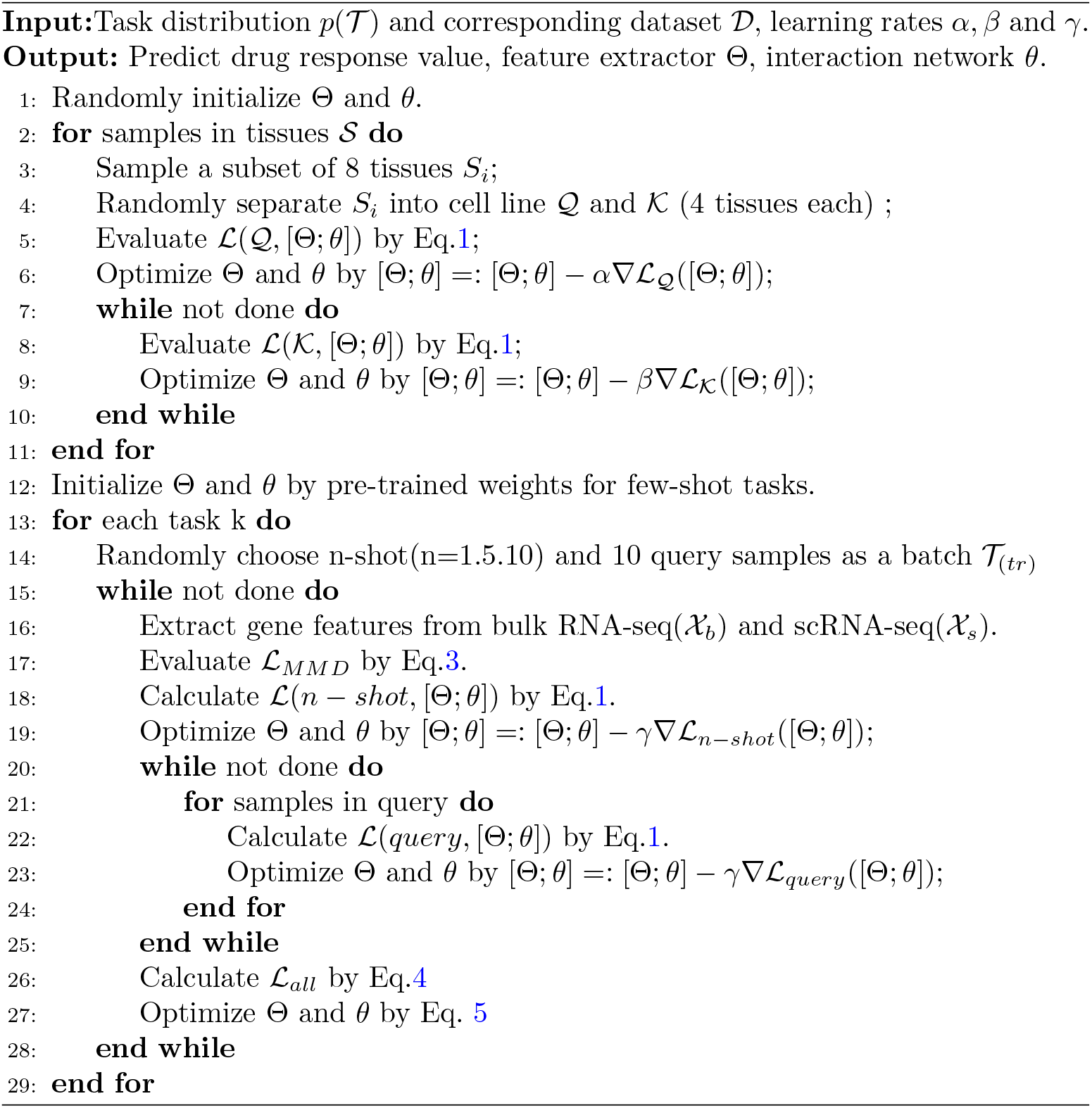

To prevent overfitting and improve training efficiency, the Dropout[36] layer is inserted after each convolutional layer. The ReLU function[37] is employed as a nonlinear activation function following each linear transformation. All models are optimized with the Adam optimizer, with the learning rate managed by StepLR. The initial learning rate is set to 0.0001 and the update step size is set to 100 iterations. The model environment is based on Python 3.10, running on the Pytorch framework of the GPU version.

#### 3.2.6 Response prediction of unseen drugs

In particular, there are scenarios where it becomes necessary to make predictions for new drugs that are not included in the training set. In this section, the performance of our model on drugs that were not part of the pre-trained GDSC dataset is explored.

This section uses the few-shot learning approach described in Section *Few-shot transfer design*, and tests on sc datasets for both the bulk-level pre-trained model and a model initialized randomly. This allows us to assess the ability of our network to adapt to new pharmacological challenges.

#### 3.2.7 Interpretability and Visualization analysis

The integral gradient (IG)[38] approach is applied to quantify the contribution of input gene features to the output predictions. An IG score is calculated for each gene by integrating the gradient of the model’s output with respect to the gene expression input, moving from a baseline of zero expression to the actual input expression level. The integration is approximated by a Riemann sum, which involves dividing the path from the baseline expression level to the input expression level into small segments and summing the gradient values across these segments:

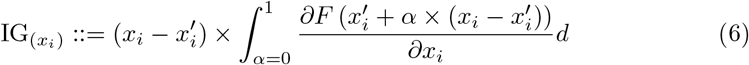

The importance of the *i*-th gene expression in the input cell *x* is calculated using a scaling coefficient *α*. The baseline expression level for gene *i*, denoted as 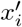, is set to zero. The notation ∂*F* (*x*_*i*_)*/*∂*x*_*i*_ refers to the gradient of the function *F* (*x*_*i*_) with respect to the *i*-th dimension, indicating how changes in the *i*-th gene expression affect the output of *F* (*x*). The input gene features is divided into 50 segments (*α* = 1*/*50) and input into the network successively to calculate the gradient. The output of the result is an IG matrix with the same shape as the input gene expression matrix. The rows represent genes and the columns represent cells. Values are IG scores. The Wilcoxon test in SCANPY is utilized to compare the integral gradient (IG) scores between the sensitive and the resistant cells, aiming to identify significantly difference genes. Genes are selected based on several criteria: a Bonferroni-adjusted *p*-value of less than 0.05, a log-fold change greater than 0.1, and the condition that more than 20% of cells in either group (sensitive or resistant) exhibited an IG score above 0.2. These selected genes are referred to as Critical Genes (CGs).

### 3.3 Model evaluation

#### 3.3.1 Evaluation metric

In this work, a regression model is constructed for the bulk dataset and a classification model for the sc dataset. For the regression model, the performance of the pre-training network was evaluated by five methods, including *R*^2^ (coefficient of determination), root mean square error (RMSE), mean square error (MSE), Spearman correlation coefficient (SCC), and Pearson correlation coefficient (PCC). For the classification model, this paper uses AUC and Average Percision (AP) to evaluate the performance on the scRNA-seq dataset, because these two evaluation indicators are less affected by the imbalanced data. Where AUC is the area under the receiver operating characteristic (ROC) curve and AP is the area under the precision-recall (PR) curve.

#### 3.3.2 Previous work

To evaluate the performance of the SCREP model, this paper compares our method with the state-of-the-art baseline models, including the results of previous work and the other deep transfer learning models.

**scDEAL**[10] is a deep transfer learning framework designed to predict cancer drug responses at the cellular level by leveraging large-scale bulk cell line data. But to the best of our knowledge, this relies on training specific model on single drug-cell line data, which poses challenges in generalizing the findings to other drugs, particularly for predicting previously unobserved drugs. At the same time, it often requires a large sample of individual cells for distribution alignment.

**SCAD**[11] is a transfer learning model designed to infer drug sensitivities at the cellular level by leveraging bulk transcriptome profiles from large public datasets of cell lines, offering insights into drug resistance and potential drug combination strategies through the identification of drug sensitivity biomarkers. However, it suffers from the same problems as scDEAL.

**ADDA**[16] is a transfer learning method first implemented on natural images. The goal of ADDA is to enable a model trained on a source domain to perform well on a target domain, even when the distribution is significantly different between the two domains. It trains a feature extractor and an adversarial discriminator to maximize and minimize the discriminability of features, respectively.

**DaNN**[17] refers to a transfer learning model designed to improve model performance on a target domain by leveraging data from a related source domain. The core idea behind DaNN is to minimize domain discrepancy and enhance generalization ability across different but related domains.

## 4 Result

### 4.1 Data statistics and visualization

For the in vitro bulk dataset, this paper used 191,034 drug-cell line pairs with normalized continuous IC50 values as labels, including 1018 cell lines with 223 drugs. This was due to the absence of response data for some drug-cell line combinations. The dataset totally covers 14 tissues and 58 tissue subtypes (see Figure 1(c) and Figure S1). The normalized distribution of total gene counts in bulk cell lines and single cells (e.g. dataset GSE110894) can be observed in Figures 2(b) and (c). The result shows that the gene expression levels in scRNA-seq are significantly lower than those in bulk RNA-seq. This disparity is likely due to differences in sample size, sequencing techniques, and underlying biological factors, which can influence the capture and detection of RNA molecules. This paper further investigated the dataset’s heterogeneity in the feature space. The UMAP of preprocessed bulk RNA-seq and all scRNA-seq (including prior-treatment and post-treatment cell populations) were plotted in Figure 2(a). This visualization demonstrates the overlap between the distributions of bulk RNA-seq and scRNA-seq in feature space. Figure 2(d) plots the UMAP visualizations of ground-truth drug response labels (sensitivity represents as 1 and resistance represents as 0), the predicted binary drug response label and the predicted continuous values of the model on the dataset GSE110894, which illustrates that our model can well distinguish the heterogeneity of drug responses.

**Fig. 2.**
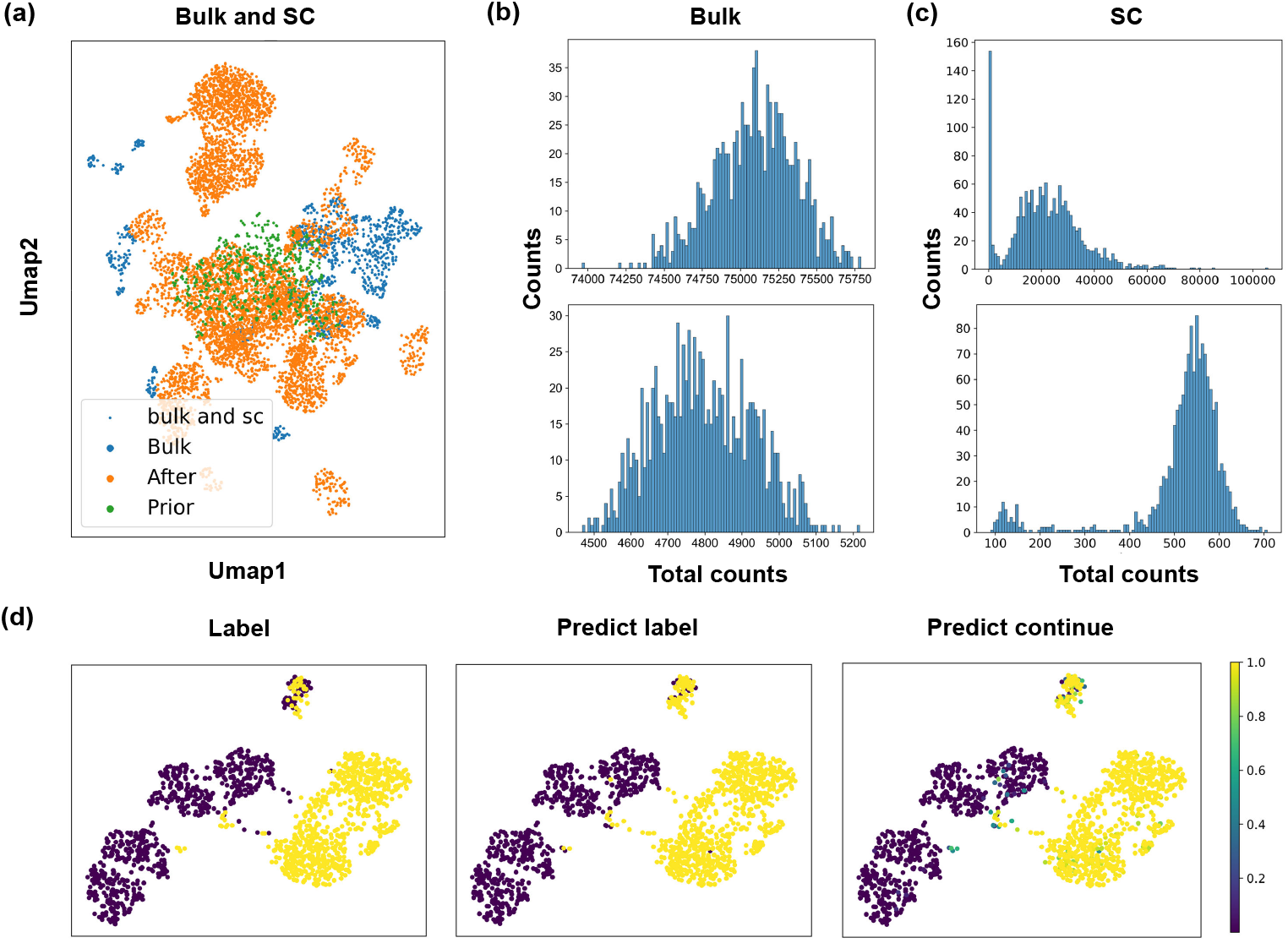
Description of the data distribution. Despite the large disparity between the bulk and SC-distributions, our model is able to transfer knowledge and distinguish well the heterogeneity of the sc response. **a**. The UMAP of preprocessed bulk RNA-seq and all scRNA-seq (including prior-treatment (prior) and post-treatment cell populations) demonstrates the overlap between the distributions of bulk RNA-seq and scRNA-seq in feature space. **b, c**. The normalized distribution of total gene counts in bulk cell lines and single cells (e.g. dataset GSE110894) before (up) and after (bottom) preprocessing shows that the gene expression in scRNA-seq was much lower compared to bulk RNA-seq. **d**. The ground-truth drug response labels (sensitivity represented as 1 and resistance represented as 0), predict binary labels and the predicted continuous values of the model.

**Fig. 3.**
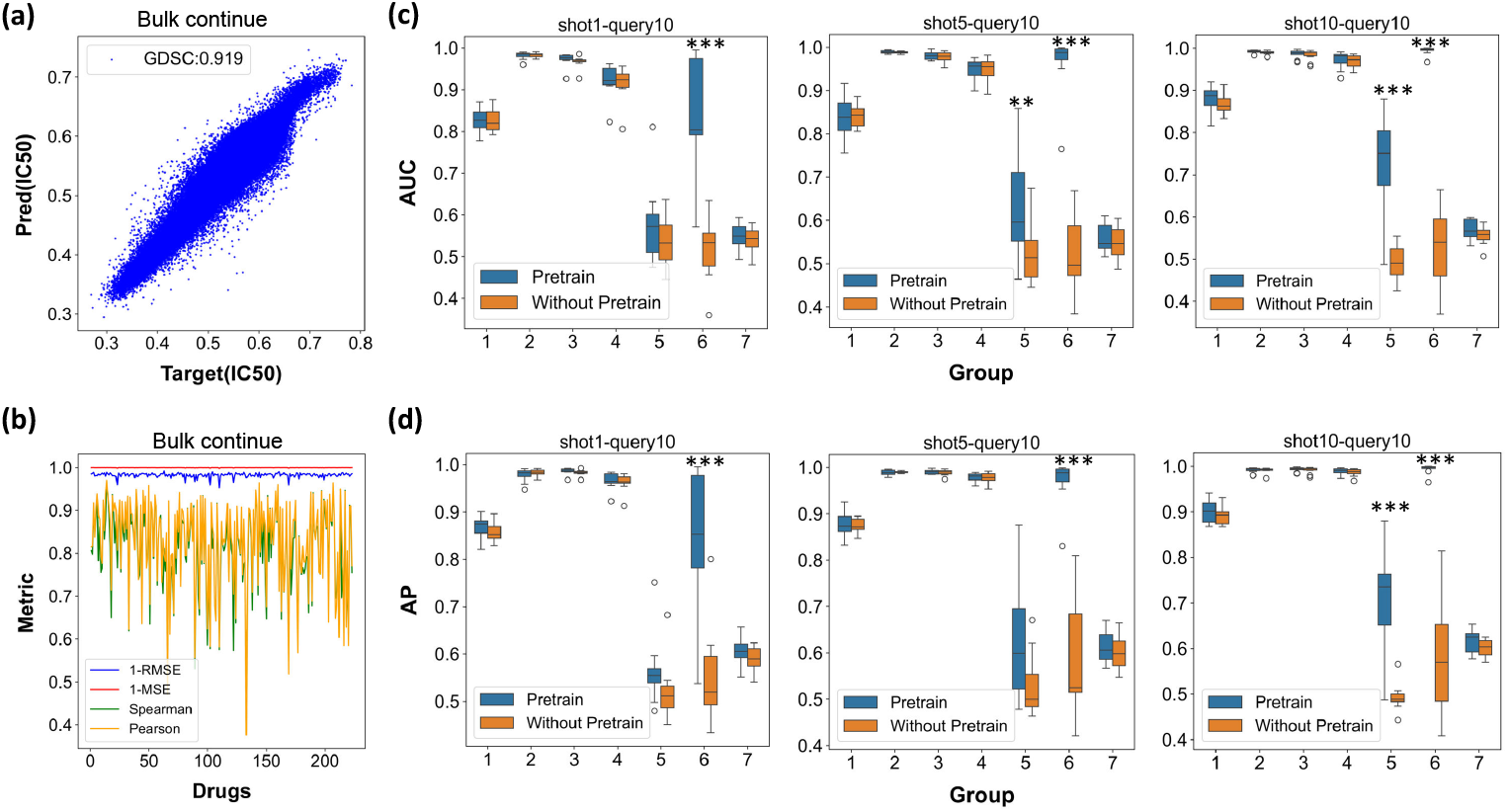
Assessment of pre-training and few-shot transfer. Our model demonstrates effective pre-training on bulk data and enhances generalization in predicting drug responses at the sc level. **a**. Linear scatter plots for the bulk dataset in the regression model. The closer the scatter is to the diagonal, the closer the model prediction is to the true value. The approximation of the model prediction to the true value achieves 0.919 *R*^2^ score. **b**. Four metrics were computed for each drug on meta pre-trained bulk data (from 1 ∼ 223), including 1-RMSE, 1-MSE, Spearman and Pearson correlation coefficient. **c, d**. Box plots of the performance of SCREP and the baseline on sc datasets with *n*-shot (*n* = 1, 5, 10) and 10-query samples transfer learning across 10 repetitions. ^*′*^*^*′*^ indicates the degree of significance of the difference. ^*′*^°^*′*^ denotes the outliers.

### 4.2 In vitro prediction model evaluation of drug sensitivity

#### 4.2.1 Meta pre-training assessment

Initially, to assess the reliability of our pre-trained model, the data is splitted into training, validation, and test sets in a 9:1:1 ratio. After completing 300 training iterations, the PCC score achieved a value of 0.940, while the SCC score reached 0.917 on the test set.

Subsequently, in the meta pre-training stage, the model’s ability to predict continuous drug responses across all bulk datasets is evaluated. SCREP achieved an *R*^2^ score of 0.919 and a PCC score of 0.959 between the predictions and the ground truth. This performance surpasses advanced deep learning models, which recorded an *R*^2^ score of 0.863 and a PCC of 0.929[39], demonstrating SCREP’s robust performance in drug response prediction for regression tasks. As shown in Figure 3(b), the four metrics calculated for each drug indicate that the response prediction values are significantly correlated with the truth, with 87.4% of them achieving a PCC value above 0.8. To investigate the impact of transfer learning, this section carries out an ablation experiment to assess model performance with and without pre-trained weights. Surprisingly, the pre-trained model significantly outperformed the non-pretrained model on prior-treatment datasets. However, on the post-treatment dataset, pre-trained model only resulted in a minor improvement. Furthermore, this paper futher compares our findings with those of previous studies, as detailed in Table 2. In contrast to previous studies, SCREP shows its competitive ability to distinguish subpopulation response heterogeneity.

**Table 2.**
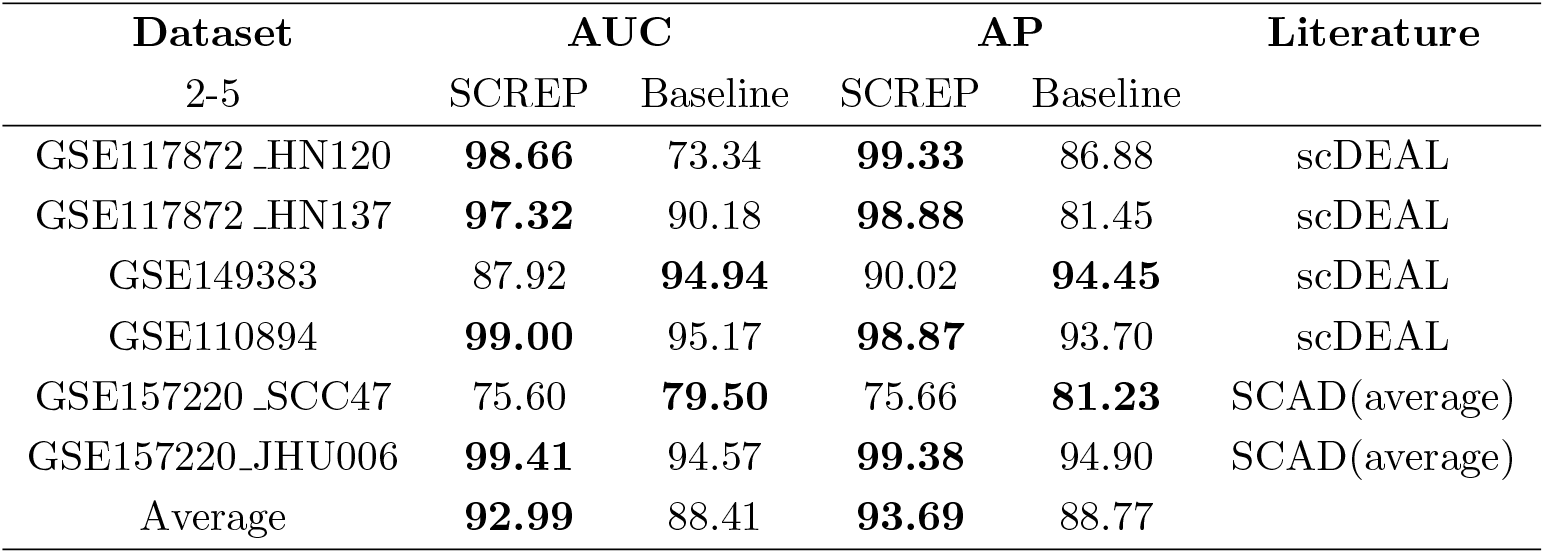
Comparison of SCREP to previous studies on sc datasets. In contrast to previous studies, SCREP shows its competitive ability to distinguish cell subpopulation response heterogeneity.

However, it is essential to recognize that bulk-based transfer learning is not a universal solution. Taking the GSE117872 dataset as an illustration (refer to Supplementary Figure S2), substantial variations can be observed in the feature space between the same patient’s cell line before and after drug intervention. Our insights from bulk RNA-seq primarily focus on the “pri” feature space, which is about the transcriptome information from pre-existing resistant cell lines. The limited improvement after transitioning to the post-treatment sc dataset is most likely due to the disparity in the feature space of gene expression before and after drug selection. Specifically, bulk data lacks transcriptomic information under drug selection, and thus knowledge transfer remains limited.

#### 4.2.2 Few-shot transfer assessment

This paper further compare the few-shot transfer strategy with other transfer learning methods, including Adversarial Discriminative Domain Adaptation (ADDA) and Domain-adaptive Neural Network (DaNN). These methods, widely used in computer vision, focusing on transferring knowledge from the source domain to the target domain. We benchmark various transfer learning methods to evaluate their applicability within our proposed integrated multimodal framework. The experimental results, detailed in Table 3, indicating that domain adaptation and adversarial learning methods are less effective at implementing transfer from the source domain to the target domain compared to the SCREP approach. This is probably due to the significant difference between the two sequencing techniques. However, this challenge could potentially be mitigated by incorporating few-shot labels, which could help bridge the gap between source and target domain in these complex transfer scenarios.

**Table 3.** Comparison of SCREP to other deep transfer learning methods on sc datasets. It indicates that domain adaptation and adversarial learning methods are less effective at implementing transfer from the source domain to the target domain compared to the SCREP approach.

| Dataset<br>2-7 | AUC |  |  | AP |  |  |
| --- | --- | --- | --- | --- | --- | --- |
|  | SCREP | ADDA | DaNN | SCREP | ADDA | DaNN |
| GSE117872_HN120 | **98.66**±0.92 | 28.63±4.39 | 52.13±13.93 | **99.33**±0.46 | 55.27±0.25 | 69.10±9.51 |
| GSE117872_HN137 | **97.32**±1.73 | 55.69±3.59 | 51.24±14.71 | **98.88**±0.68 | 74.35±0.23 | 72.32±8.76 |
| GSE149383 | **87.92**±2.84 | 50.65±0.24 | 48.48±4.23 | **90.02**±1.99 | 59.99±0.12 | 43.3±3.22 |
| GSE110894 | **99.00**±0.87 | 38.85±1.03 | 54.51±10.65 | **98.87**±1.27 | 46.79±0.72 | 58.01±8.63 |
| GSE157220_SCC47 | **75.60**±14.07 | 56.99±1.53 | 50.13±1.09 | **75.66**±14.00 | 52.96±0.18 | 50.64±1.37 |
| GSE157220_JHU006 | **99.41**±0.78 | 47.28±2.29 | 50.47±3.45 | **99.38**±0.87 | 41.06±0.98 | 50.81±2.25 |
| GSE228154 | **57.03**±2.33 | 47.73±0.13 | 49.86±1.03 | **61.64**±2.33 | 55.48±0.09 | 55.52±1.03 |

### 4.3 Unseen drug response prediction assessment

The transferability of gene expression-molecular interaction knowledge is evaluated using a prior-treatment sc dataset that includes drugs not present in the GDSC dataset. Specifically, SCREP is tested on the remaining 108 drugs in GSE157220. This compares the adaptation capabilities under both pre-trained and non-pre-trained conditions after 100 iterations. Since a dynamic threshold may not be feasible in real-world applications, a fixed threshold of 0.5 is set for both training and testing. F1 score is used to comprehensively evaluate the performance of the network. As a result, the pre-trained group achieved an average F1 score of 87.06%, compared to 67.03% for the non-pre-trained group, showing about a 20% improvement. Examples of drugs are listed in Table 4, and results for all the 108 drugs are provided in Supplementary Data S1.

**Table 4.** Unseen drug response prediction on w/o pre-trained and pre-trained model. Examples of drugs are listed below, and results for all 108 drugs are provided in Supplementary Data S1.

| Unseen drug in<br>GDSC | w/o pre-trained<br>(shot5) | pre-trained (shot5) |
| --- | --- | --- |
| PRT062607 | 81.20 | **100.00** |
| BAY 11-7082 | 38.19 | **89.69** |
| ZSTK474 | 33.33 | **97.73** |
| Vorinostat | 58.09 | **100.00** |
| YM201636 | 38.19 | **97.73** |
| Cediranib | 67.63 | **93.15** |

### 4.4 Case Study: Identify critical genes and pathways for drug response

While SCREP achieves excellent results in predicting drug responses at cellular level, it is crucial to determine whether the model prediction results are reliable in clinical scenarios. The aim is to explore the correlation between drug response and gene expression in order to explore pathways of drug action, evaluate the validity and reliability of the model and provide new methods for the discovery of possible targets. This section conducts a case study for oral squamous cell carcinoma (OSCC) treated with Cisplatin (dataset GSE117872 HN120). Cisplatin acts by generating DNA crosslinks via interactions with purine bases, causing DNA damage and interfering with DNA replication, leading to cell apoptosis, particularly in rapidly dividing cells such as cancer cells. Consequently, cells sensitive to Cisplatin may express genes associated with apoptosis[40].

The results shown that cells showing resistance may have large distance in the feature space, as shown in Figure 4(a) for subgroups A and B. Nonetheless, SCREP is able to effectively distinguish resistance in cells with similar cluster spaces (subgroups A). The model achieves an AUC of 0.99, with only a few predictions failing, as indicated in Table 3 and Figure 4(a). Critical genes are filtered based on p-values *<*0.05, log-fold changes *<*0.1, and more than 20% of cells in either comparison group exhibited a significant expression difference. The results show that of all 16,906 genes, 1,897 are up-regulated in drug-sensitive cells and 1,712 are up-regulated in drug-resistant cells, both showing significantly different attribution gradients. Now, considering both up-regulated and down-regulated genes in the resistant group compared to the sensitive group, genes with a log-fold change greater than threshold *th*1 are considered to be significantly up-regulated, while those with a log-fold change lower than threshold *th*2 are considered to be significantly down-regulated (Figure 4(b)). The distribution of the log-fold changes is analyzed for genes and determined thresholds *th*1 and *th*2 at the 5% and 95% percentile, respectively. The top 5 up-regulated and down-regulated genes are selected and plotted on a violin graph (Figure 4(c)). Gene set enrichment analysis (GSEA) is implemented on the 1897 drug-sensitive genes and 1712 drug-resistant genes by Decoupler[41]. The significant enrichment pathways are filtered by p-values *<*0.05, and the results are presented in Figure S3, S4, and Supplementary Data S2, S3. The results indicate that the genes sensitive to Cisplatin are significantly down-regulated on pathway “recognition of DNA damage by PCNA containing replication complex” (score=-0.416, p-value=0.038). Inefficient identification of DNA damage may lead to a delay or failure in the initiation of the DNA repair process, resulting in the accumulation of various DNA lesions such as mismatches, insertions, deletions and crosslinks[42, 43], which is one of the biological processes accociated with Cisplatin-sensitive. At the same time, the “DNA replication” pathway is up-regulated in drug-resistant cells (score=0.405, p-value=0.078). This enhanced replicate capacity enables cancer cells to repair Cisplatin-induced DNA cross-links more effectively, contributing to resistance.

**Fig. 4.**
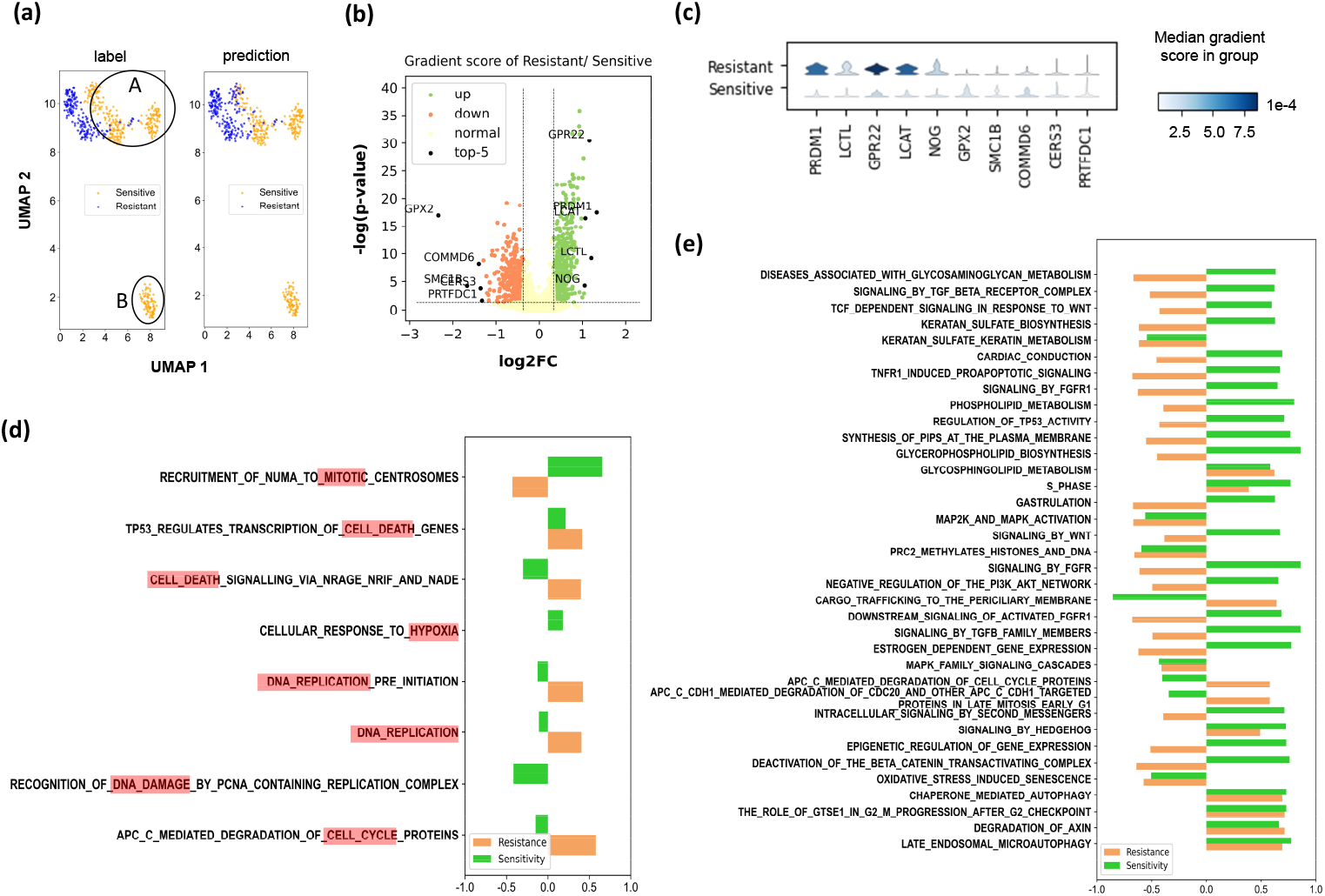
Case study of identifying critical genes and pathways for drug response on GSE117872 HN120. It demonstrates that SCREP can effectively identify critical genes and pathways associated with drug response, highlighting its utility in uncovering the complex biological dynamics that contribute to drug resistance.**a**. The UMAP of ground-truth (left) and predicted binary drug response (right). A and B are the different subpopulations with the same drug resistance label. **b**. The volcano map showed that the drug-resistant group significantly up-regulated (green) and down-regulated (orange) critical genes compared with the drug-sensitive group. **c**. The top 5 up- and down-regulated genes are plotted on a violin diagram. The color of each violin indicates the median gene gradient score of the resistant or sensitive groups. **d**. The pathways about cell mitotic and death, showing different or diametrically opposite correlations between the sensitivity and the resistance. **e**. 36 significant pathways of drug-resistant (p*<*0.05) and the correlation coefficient. Among them, the direction of regulation for resistance and sensitivity is opposite in 20 pathways.

On the other hand, it is also noted that Cisplatin-resistant critical genes are significantly enriched in the pathway related to “APC-C mediated degradation of cell cycle proteins” (score=0.579, p-value=0.019), which plays a critical role during mitosis. The Anaphase-Promoting Complex or Cyclosome (APC/C) is an E3 ubiquitin ligase that regulates the cell cycle by targeting specific cell cycle proteins for degradation, thus controlling progression through mitosis and the G1 phase of the cell cycle. The effectiveness of Cisplatin largely depends on its ability to induce DNA damage that leads to cell cycle arrest and apoptosis[29]. The up-regulation of APC/C activity can lead to an accelerated transition to mitosis through the timely degradation of key cell cycle inhibitors. This rapid progression may reduce the time available for Cisplatin-induced DNA damage to be detected and repaired before cell division. This scenario potentially allows DNA-damaged cells to continue dividing, which can contribute to the development of resistance.

Specifically, 36 significant pathways are identified to be associated with drug resistance (p*<*0.05), as illustrated in Figure 4 (e). In this analysis, the correlation coefficient measures the relationship between the drug response predicted by SCREP and each pathway. A positive enrichment score indicates that a pathway is up-regulated or enriched in the population, whereas a negative enrichment score suggests that the pathway is down-regulated or depleted. Remarkably, the direction of regulation is opposite in 20 pathways compared to the drug-sensitive group. Additionally, other critical pathways, such as those involved in cell mitosis and death, are listed in Figure 4 (d). These pathways show different or even diametrically opposite correlations between sensitivity and resistance. This analysis demonstrates that SCREP can effectively identify critical genes and pathways associated with drug response, highlighting its utility in uncovering the complex biological dynamics that contribute to drug resistance.

### 4.5 Generalization on other cell line sequencing platforms

This section repeated experiments on the CCLE dataset, pre-trained the model from scratch, and validated it on the single-cell dataset. The results are shown in Figure S5 and Figure S6. The results shown that CCLE generally generalizes better on single-cell datasets than the results without pre-training.

### 4.6 Parameter sensitivity analysis

In the section on experimental results (Figure 3(c) and (d)), the outcomes of utilizing various numbers of samples in the support set have compared. In this part, we investigate the effects of employing different quantities of samples as query sets on the performance of few-shot learning. The results are shown in the supplementary of Figure S1. It is shown that both too large and too small queries affect the capabilities of the model. In the experiments, query=10 was chosen based on the total number of samples and the performance of the model.

## 5 Discussion

The analysis of single-cell pharmacogenomics elucidates the resistance mechanisms present within specific cell populations, which is advantageous for precision medicine. However, these resistance mechanisms are highly complex and involve numerous factors. While pre-training with data from cancer cell lines cultured in vitro provides a foundational basis, transferring this knowledge to single-cell contexts still poses significant challenges. We infer that these challenges primarily arise from alterations in the post-treatment transcriptome and the availability of transferable multimodal data.

### Changes in the post-treatment transcriptome

Pre-clinical models developed using pre-treatment transcriptome data can be used for drug screening; However, they may struggle to accurately predict drug efficacy at a later stage when relying on post-treatment transcriptome data. The underlying reason for this may be an insufficient understanding of the bulk transcriptome following the treatment, which requires significant engagement of biochemists and considerable accumulation of experimental data.

### Multimodal data transfer learning

Our framework is constrained by the uni-modal gene expression information available in the sc dataset. This limitation hinders the development of pre-trained models that can leverage multiple modalities, such as genetic mutations and methylation patterns. Expanding the model to incorporate these data types may significantly enhance its prediction accuracy and allow for a more comprehensive understanding of drug responses at the cellular level.

## 6 Conclusion

In summary, this paper proposes SCREP to predict cellular responses to multiple drugs, which overcomes the limitations of model generalization. Firstly, a meta pretraining and few-shot adaptive distribution alignment strategy are implemented for cross-domain transfer while minimizing the sample size requirement. Secondly, the proposed model can generalize to drugs not encountered during pre-training, suggesting that the model can be extended to a wider range of sc-based drug screening applications. Thirdly, attribution gradient methods can be employed to identify crucial genes and pathways under multi-modal inputs, providing a new way to understand the resistance mechanisms. Our work could potentially guide clinical precision therapy by predicting resistance based on scRNA-seq data. In the long term, SCREP may also benefit the virtual screening of anti-cancer drugs discovery.

## Supplementary Materials

**Supplementary material S1**. Figure S1∼S6

**Supplementary Data S1**. Results of all unseen drug response prediction evaluations.

**Supplementary Data S2**. GSEA results of the 1897 drug-sensitive genes.

**Supplementary Data S3**. GSEA results of the 1712 drug-resistant genes.

## Data availability

All datasets analyzed in this study are publicly available. Details of the data resources are described in Section Materials and methods. The drug response information can be downloaded from the page GDSC download and CCLE PRISM download. The cell lines of GDSC is publicly available through the website GDSC1000 WebResources. The cell lines of CCLE is publicly available through the websiteCCLE bulk download. The origion single cell dataset can be found on GEO by GEO identifiers or origional paper: GSE117872, GSE149383, GSE110894, GSE157220, GSE228154. The genetic symbols from PanglaoDB dataset can be obtained from PanglaoDB. The preprocessed single-cell data set is available for download by following the instructions in the code link.

## Code availability

The code used to train and generate results can be found at https://github.com/geshuang307/SCREP.

## Author contributions

**Shuang Ge:** Conceptualization, Data curation, Formal analysis, Investigation, Methodology, Software, Validation, Visualization, Writing – original draft, Writing – review & editing. **Shuqing Sun:** Writing – original draft, Writing – review & editing. **Yiming Ren:** Writing – original draft, Writing – review & editing. **Huan Xu:** Methodology, Writing – original draft, Writing – review & editing. **Qiang Cheng:** Supervision, Resources, Conceptualization, Writing – original draft. **Zhixiang Ren:** Conceptualization, Formal analysis, Funding acquisition, Methodology, Project administration, Resources, Supervision, Writing – original draft, Writing – review & editing.

## Acknowledgements

We appreciate the Peng Cheng Cloud-Brain.

## Footnotes

1 https://depmap.org/portal/datapage/?tab=allData

2 https://www.ncbi.nlm.nih.gov/geo/

3 https://PanglaoDBdb.se/

4 https://www.ncbi.nlm.nih.gov/gene

