## Supplementary Figure S1-S4 for "SCREP: Towards Single-Cell Drug Response Prediction by Pharmacogenomic Embedding Enhanced Meta-Pretraining and Few-Shot Transfer Learning" (from another version of this preprint: version 1, 2024-04-28)

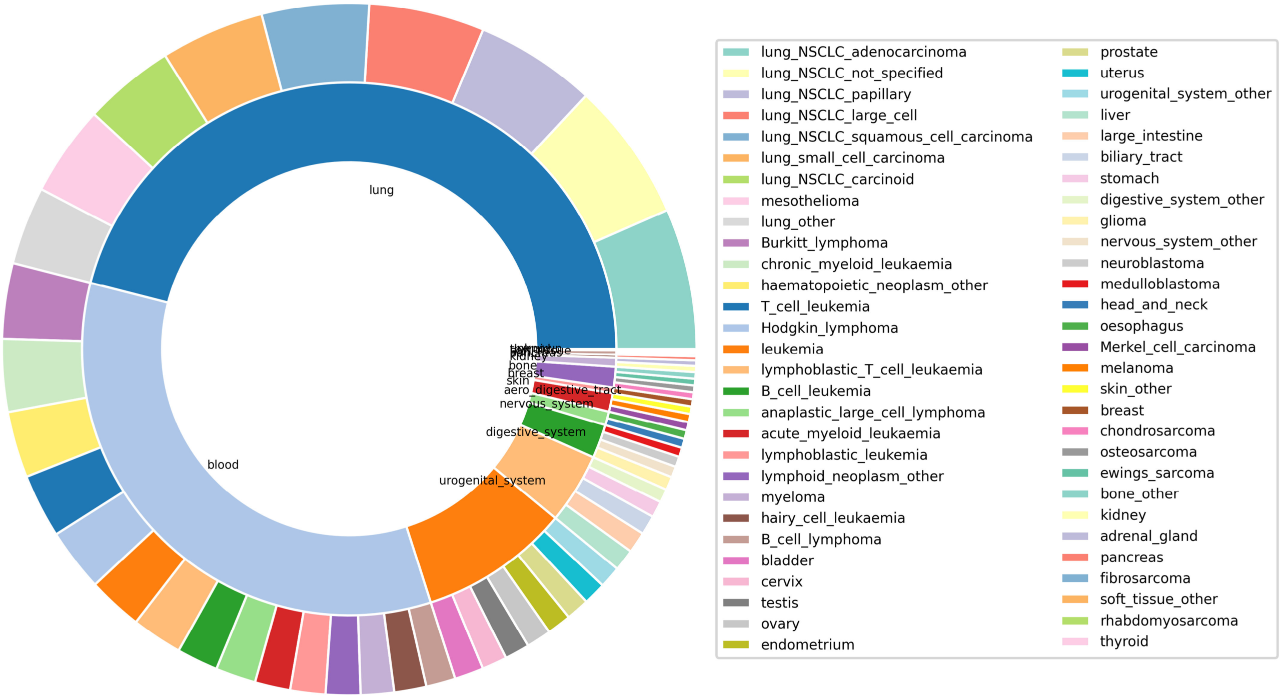


**Figure 1: Supplementary Figure S1.** Dataset statistic shows the distribution of bulk datasets, including 14 tissues with 58 subtissues.

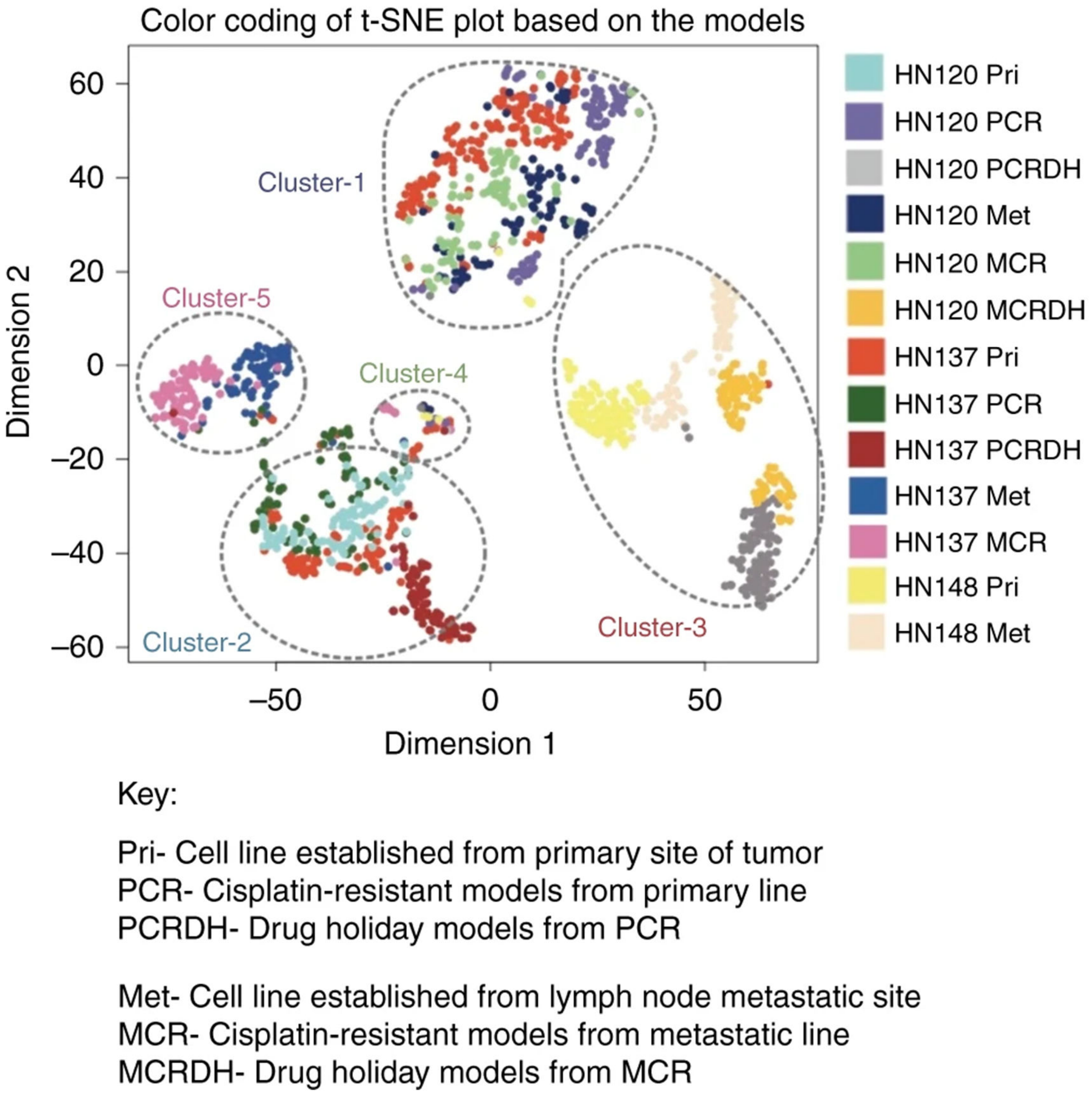


**Figure 2: Supplementary Figure S2.** Color-coding of t-SNE plot based on the identities of individual OSCC-PDPCs and their drug-resistant/holiday models. In Fig. 2d of the original paper[1].

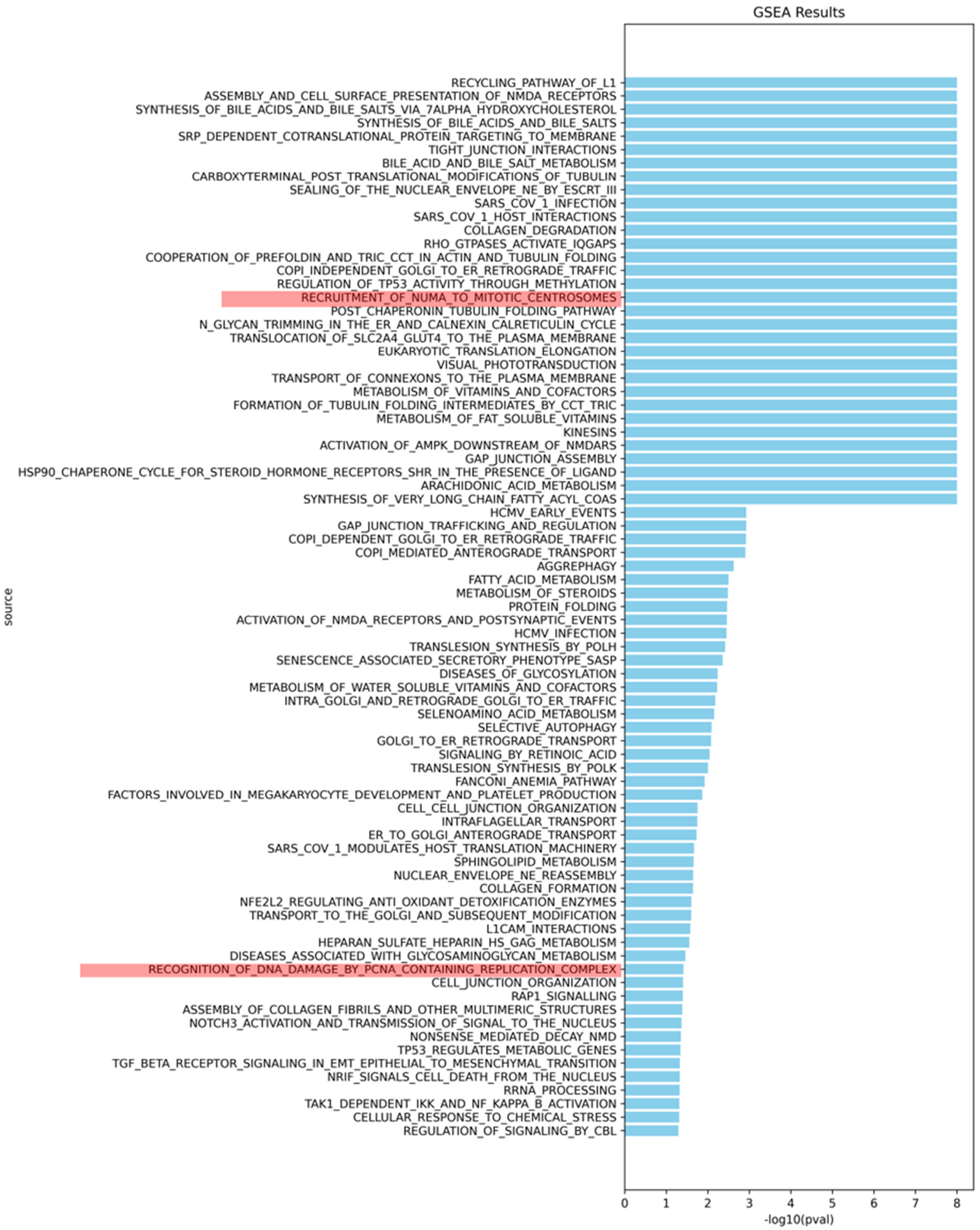


**Figure 3: Supplementary Figure S3.** Gene set enrichment analysis (GSEA) of the 1897 drug-sensitive genes, the result shows 79 significant enrich pathways.

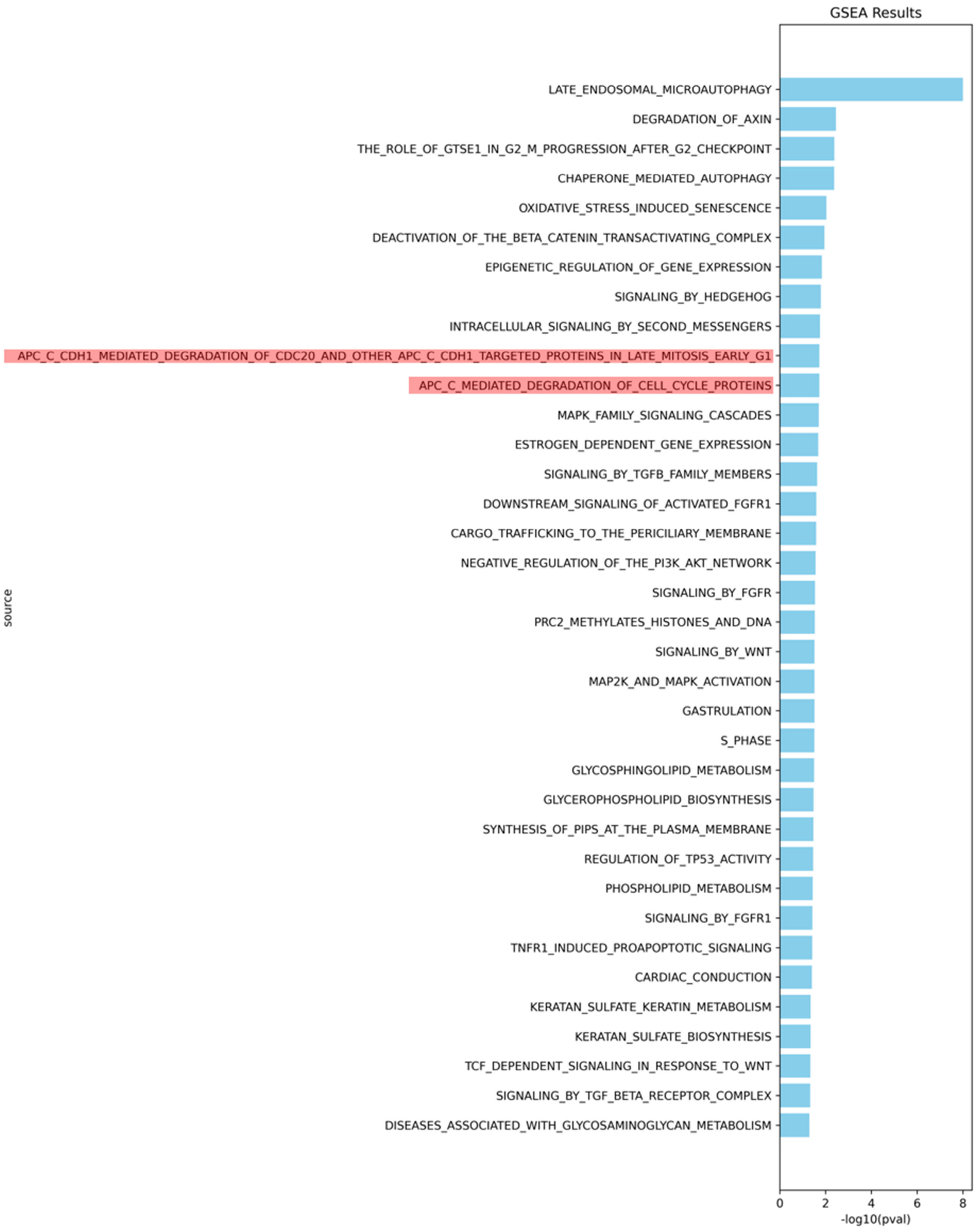


**Figure 4: Supplementary Figure S4.** Gene set enrichment analysis (GSEA) of the 1712 drug-resistant genes, the result shows 36 significant enrich pathways.
